# Functional and molecular rescue of aganglionic colon by human enteric nervous system progenitor transplantation in Hirschsprung disease

**DOI:** 10.1101/2025.10.16.679985

**Authors:** Benjamin Jevans, Fay Cooper, Luca Peruzza, Silvia Perin, Shuqi Li, Tara McAteer, Paolo De Coppi, Peter W Andrews, Anestis Tsakiridis, Conor J McCann

**Affiliations:** Stem Cells and Regenerative Medicine, UCL Great Ormond Street Institute of Child Health, London, UK; NIHR Great Ormond Street Hospital Biomedical Research Centre, London, UK; School of Biosciences, The University of Sheffield, Sheffield, UK; Department of Comparative Biomedicine and Food Science, University of Padova, Padova, Italy; Department of Woman and Child Health, University of Padova, Padova, Italy; Specialist Neonatal and Paediatric Surgery Unit, Great Ormond Street Hospital, London, UK

## Abstract

Hirschsprung disease (HSCR) is a devastating congenital disorder characterised by absence of the enteric nervous system (ENS) in the distal gut. Cell therapy, using human pluripotent stem cell-derived ENS progenitors, offers an attractive alternative to treat this life-limiting disorder. Here we provide an in-depth characterisation of the HSCR phenotype in the B6;129-*Ednrb^tm1Ywa^*/J mouse model including molecular analyses highlighting the wide-ranging molecular effects of aganglionosis in HSCR colon. We show that transplantation of hPSC-derived ENS progenitors leads to significant increases in contractile function with formation of extensive donor-derived neuronal networks in colonic explants. Moreover, post-transplantation, we show that integration of ENS progenitors rescues many of the biological pathways impacted in the HSCR aganglionic microenvironment. These novel results provide strong evidence that human ENS progenitor transplantation can modulate critical signalling pathways affected in HSCR and exert positive impacts on contractile function, in aganglionic tissue, supporting further translational development of this regenerative medicine approach.

## Introduction

Approximately 1 in every 5000 live births are affected by Hirschsprung disease (HSCR) (1). The disease affects the enteric nervous system (ENS), the intrinsic innervation of the gastrointestinal (GI) tract (2). The ENS is responsible for multiple functions in the GI system including control of gastric digestion, nutrient absorption, monitoring the intestinal lumen environment and gastrointestinal motility to propel digesting food along the tract. In the absence of an ENS, the musculature of the GI tract tonically constricts, leading to an accumulation of intestinal contents in the proximal, ganglionic colon and the development of megacolon (3). Without treatment interventions, this distended region is prone to Hirschsprung disease-associated enterocolitis (HAEC) and rupture: potentially life-threatening situations (4). Surgery is lifesaving, involving resection of the aganglionic bowel and a ‘pull through’ of the ganglionic bowel into a necessarily retained portion of distal, aganglionic anus (5). However, patients often require multiple follow up surgeries and can suffer lifelong complications, including recurrent enterocolitis, chronic abdominal pain, constipation and faecal incontinence (6). Hence, further research into alternative treatments is urgently required.

The neuroglial cells of the ENS are derived predominantly from vagal neural crest cells (NCCs), which arise at the neural tube, migrate to the oral end of the developing foregut and proceed to colonise the entire GI tract in an oro-anal direction (7, 8). In HSCR, however, this colonization is severely impaired, leaving a variable portion of the distal colon (and occasionally even the entire colon and parts of the small intestine) aganglionic (9). The causes of this appear to be multifactorial and several genes and signalling pathways have been identified as risk factors (10). For example, *Ednrb* (Endothelin receptor b) and its ligand *Edn3* (Endothelin 3) are believed to be vital for correct colonization of the gut by NCCs (11–13) and mutations in this signalling pathway have been shown to result in human HSCR (14).

Stem cell therapy offers the potential to replenish missing neuroglial cells in the aganglionic colon and restore correct gastrointestinal motility. The recent development of protocols allowing for rapid generation of human pluripotent stem cell (hPSC)-derived ENS progenitors may provide an ideal source of cells to compensate for the failed colonization observed in HSCR (15). Indeed, human ENS progenitors generated using these protocols have recently been shown to significantly increase the contractile phenotype of human HSCR surgical discard tissue explants following *ex vivo* transplantation (16). However, while preliminary investigations have provided insights into the mechanisms of donor ENS cell integration in gut tissue (17), an assessment of the full molecular impact of donor ENS progenitor integration post-transplantation, in recipient tissue, is lacking. Human surgical discard tissue samples are rare and heterogeneous, limiting study of the molecular changes induced in the aganglionic microenvironment by donor cells. Several groups have used various transgenic HSCR mouse models to demonstrate survival and integration of transplanted cells as well as functional improvements (18–20). These models, including mouse lines targeting the *Ednrb* gene (11), provide an invaluable tool for studying the development and progression of HSCR, as well as for testing novel therapies such as cell-based interventions. However, phenotypic variability between differing strains targeting the Ednrb gene has hampered efforts to compare studies using different transgenic models. Indeed, an in-depth molecular characterization of the aganglionic microenvironment in many of these models has not been performed and an understanding of the impact of cell transplantation to recipient colonic tissue in such models is also lacking, compounding comparison issues.

Here, to better understand the aganglionic microenvironment in HSCR and define the impact of human ENS progenitor transplantation, we provide an in-depth phenotypic characterisation of the B6;129-*Ednrb^tm1Ywa^*/J mouse model at the cellular and molecular level. Further, we employ a reproducible explant platform, similar to that used in human tissue explants (16), to test the impact of human ENS progenitor-based transplantation. Using this approach, we show significant increases in contractile activity in aganglionic colonic tissue transplanted with hPSC-derived ENS progenitors, as well as the formation of transplant-derived, ganglia-like structures. Importantly, we also demonstrate that ENS progenitor transplantation triggers *de novo* synaptogenesis within aganglionic gut segments, with donor-derived cells differentiating to numerous neuronal lineages, extensive reconfiguration of the host microenvironment with increased vasculogenesis and a potentiation of migratory characteristics within human donor cells themselves. Taken together, these results highlight the utility of the B6;129-*Ednrb^tm1Ywa^*/J mouse model as a tool for studying the molecular impact of ENS progenitor transplantation and strongly support the use of hPSC-derived ENS progenitors as a cell therapy for HSCR.

## Results

### The B6;129-*Ednrb^tm1Ywa^*/J mouse model phenocopies human Hirschsprung disease

B6;129-*Ednrb^tm1Ywa^*/J mice are known to display colonic aganglionosis and reproducibly die before breeding age (11) as opposed to some other HSCR mouse models, such as those incorporating the piebald (*Ednrb^s^)* and piebald lethal (*Ednrb^sl^)* alleles, in which significant numbers of homozygotes survive and can breed (21, 22) (https://www.jax.org/strain/000308). This reproducibility makes B6;129-*Ednrb^tm1Ywa^*/J mice an ideal model for studying the effects of cell-transplantation. However, to the best of our knowledge, the B6;129-*Ednrb^tm1Ywa^*/J mouse model has never been fully characterised. Therefore, before using this model to test the potential benefits of ENS progenitor cell transplantation, we undertook an in-depth characterization of this model.

B6;129-*Ednrb^tm1Ywa^*/J mice, hereafter denoted *Ednrb^-/-^*, were maintained on a heterozygous background, with knockout offspring easily identified by their skewbald coat colouring (**Fig. S1A**) and confirmed by genotyping (**Fig. S1B**). By postnatal day (P) 21, with rare exceptions, the abdomen of *Ednrb^-/-^* pups, became distended and their locomotion increasingly restricted, necessitating culling to remain within humane endpoints. GI tracts of wildtype (WT) *Ednrb^+/+^* littermates, isolated at P14, displayed a uniform diameter with evidence of well-formed faecal pellets in the colon (**Fig. S1C**). In stark contrast, isolated GI tracts of *Ednrb^-/-^* offspring displayed a constricted distal portion of the colon with a severely swollen ‘megacolon’ proximal to this constricted region (**Fig. S1D**). To determine whether the presence of a homozygous knockout mutation was embryonic lethal, all pups from 13 litters (derived from 4 separate breeding pairs, 124 pups in total), were genotyped to determine the relative ratio of homozygotes, heterozygotes and WT offspring (**Fig. S1E**). We found that 27.35% were *Ednrb^+/+^*, 18.8% were *Ednrb^-/-^* and 53.84% were *Ednrb^+/-^*, in line with predicted Mendelian inheritance patterns. Chi squared analysis confirmed no significant difference between these and the expected ratios (25% *Ednrb^+/+^*, 25% *Ednrb^-/-^*and 50% *Ednrb^+/-^*, p=0.3009), implying that the *Ednrb^-/-^*genotype is not embryonic lethal. A subset of *Ednrb^-/-^* colons were labelled with Phox2b to map the distribution of enteric ganglia (**Fig. 1A-D**). This revealed dense ganglia in the proximal colons of *Ednrb^-/-^* mice, formed of clustered Phox2b+ nuclei (**Fig. 1B**). The frequency of these ganglia was substantially reduced in the transition zone (**Fig. 1C**) and Phox2b+ nuclei were almost exclusively absent in the distal *Ednrb^-/-^*colon (**Fig. 1D**).

**Figure 1:**
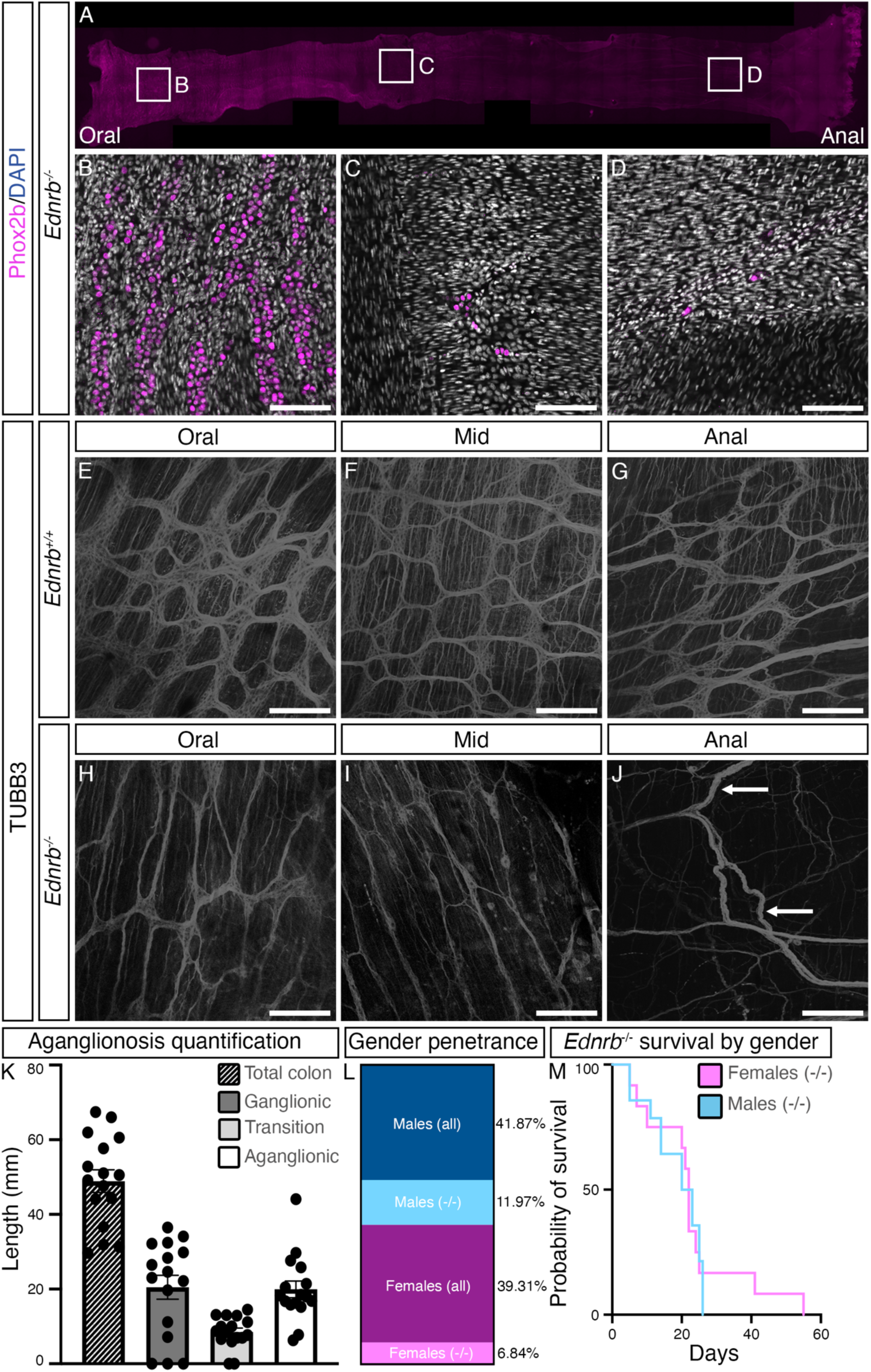
Variable severity of colonic aganglionosis in B6;129-*Ednrb^tm1Ywa^*/J mice is lethal but unaffected by gender. (A) Colonic tissue isolated from *Ednrb^-/-^* animals was stained with antibodies raised against Phox2b. (B-D) Higher magnification of Phox2b staining in the ganglionic, proximal colon (B), partially ganglionated transition zone (C) and aganglionic, distal colon (D). (E-G) TUBB3 staining of *Ednrb^+/+^* wild type littermates confirms the characteristic ganglia of the enteric nervous system throughout the colon, including proximal (E), mid (F) and distal colon (G). (H-J) Conversely, *Ednrb^-/-^* animals display ganglia in the proximal region (H), the density of which is much reduced in the mid colon (I). (J) In the distal colon ganglia are entirely absent and hypertrophied nerve bundles can be seen (J, arrows). (K) The length of the proximal, ganglionic and distal, aganglionic segments were approximately equal in *Ednrb^-/-^* animals, with a shorter transition zone. (L) Approximately twice as many males were affected as females. (M) Survival of *Ednrb^-/-^* animals was unaffected by gender. Scale bars: B-D – 100μm, E-J – 200μm. Error bars are standard error of the mean.

To further characterise the aganglionic colon, including examination of both neuronal cell bodies and extensions, muscularis tissue isolated from *Ednrb^+/+^* and *Ednrb^-/-^* animals were stained with the pan-neuronal marker TUBB3. *Ednrb^+/+^* mice displayed a stereotypical pattern of TUBB3+ ganglia in the proximal, mid and distal colon (**Fig. 1E-G**). However, while the proximal colons of *Ednrb^-/-^* mice were ganglionic, this gave way to a characteristic ‘transition zone’ of colonic tissue with a reduced density of ganglia before, in the distal colon, these ganglia were lost entirely (**Fig. 1H-J**). Notably, as has been previously reported in human HSCR tissue samples (23, 24), the aganglionic distal colons of *Ednrb^-/-^* mice often displayed substantial, hypertrophied innervation, presumably extrinsic in origin (**Fig. 1J**, arrows). We quantified the extent of colonic aganglionosis by demarcating three distinct colonic regions: ganglionic, defined as an area with a uniform distribution of ganglia; transition, defined as an area with some ganglia but noticeably less than in the ganglionic regions; and aganglionic, lacking ganglia entirely. The average total colon length was 48.93mm +/-3.032mm, of which 20.48mm was ganglionic +/-3.198mm, 8.5mm was transition zone +/-1.058mm and 19.93mm was aganglionic +/-2.228mm (**Fig. 1K**). Survival of *Ednrb^-/-^* mice was significantly, inversely correlated to the extent of aganglionosis (**Fig. S1F**, p=0.0447).

In humans, HSCR is known to exhibit a male:female gender bias of almost 4:1 (25). Across 11 litters, 53.84% of live births from *Ednrb^+/-^* breeding pairs were male and 46.15% were female. 11.97% of all pups were male *Ednrb^-/-^*, while only 6.84% of all pups were female *Ednrb^-/-^*, displaying a gender bias of approximately 2:1 in favour of males (**Fig. 1L**). However, Chi squared analysis revealed that this was not significantly different from expected ratios (12.5% male *Ednrb^-/-^*, 12.5% female *Ednrb^-/-^*, p=0.2922). The severity of colonic aganglionosis was unaffected by gender (**Fig. S1G**), with males displaying an average ganglionic tissue length of 20.16 +/-4.398mm, a transition zone tissue length of 9.25 +/-1.497mm and an aganglionic tissue length of 21.52 +/-3.084mm. Comparatively, females displayed an average ganglionic tissue length of 20.9 +/-5.023mm, a transition zone length of 7.55 +/-1.509mm and an aganglionic tissue length of 17.9 +/-3.29mm. Gender also had no effect on survival (**Fig. 1M**): the median survival was 18.79 +/-2.041 days for males and 22.83 +/-4.005 days for females (p=0.3566). As gender had no apparent effect on the disease severity, we used male and female mice in all subsequent experiments. Together, these findings reinforce the use of the B6;129-*Ednrb^tm1Ywa^*/J model as a key platform for studying the effects of variable aganglionosis and regenerative medicine approaches for HSCR.

### Key neuronal pathways are significantly downregulated in *Ednrb^-/-^* colon

To determine the molecular changes associated with an HSCR-like aganglionic phenotype, we undertook bulk RNAseq using the *tunica muscularis* from the distal 1/3 of the colon in *Ednrb^-/-^* mice and WT littermates (n=3 for each). Multi-dimensional scaling (MDS) (26) showed that the samples clustered according to genotype (**Fig. S2**). Upon further analysis, 1139 genes were found to be differentially expressed between *Ednrb^-/-^* and WT littermate tissue samples (**Fig. 2A, Table S1**). Over-representation analysis of the down-regulated genes, using the Kyoto encyclopaedia of genes and genomes (KEGG) pathway database, identified numerous neuronal pathways impacted in *Ednrb^-/-^* tissue compared to WT littermate tissue, including *Cholinergic Synapse*, *Dopaminergic Synapse*, *GABAergic Synapse*, G*lutamatergic Synapse* and *Serotonergic Synapse* (**Fig. 2B, Table S2**). To determine whether the activity of these pathways was likely to be affected by the differentially expressed genes, we used signalling pathway impact analysis (SPIA, **Fig. 2C, Table S3**), which considers the fold change of a given gene in combination with its likely impact within a pathway (i.e., activating or inhibiting) (27). This revealed a significant inhibition of the activity of the *Cholinergic Synapse* (the most severely impacted), *Dopaminergic Synapse*, *GABAergic Synapse* and *Serotonergic Synapse* pathways, with the only exception being the *Glutamatergic Synapse* pathway activity (which was likely to be increased). Since the symptoms of HSCR are believed to be driven primarily by the loss of neuronal activity, we next examined the specific genes affected in each neuronal pathway significantly downregulated in *Ednrb^-/-^* tissue compared to WT littermate tissue (**Fig. 2Di**). Genes of interest identified in this analysis included *Edn3*, the signalling partner of *Ednrb*, in addition to numerous other crucial neuronal genes, such as *Nos1* and *Snap25*. Interestingly, numerous downregulated genes in *Ednrb^-/-^* tissue were present in multiple pathways (**Fig. 2Dii**). This included ion channels/transporters such as *Kcnq5*, *Kcnn2* and *Atp2b2*; numerous proteins required for neuronal function such as *Npy*, *Unc13a*, *Nmu*, *P2rx2*, *Nos1* and *Mapk10* genes; and genes involved in satiety such as *Cckar*. These results highlight that disruption of Ednrb signalling leads to significant impacts on numerous genes and modulation of multiple biological pathways within the aganglionic microenvironment. Of note, the presence of these affected genes in multiple pathways may make them attractive targets for pharmacological interventions in the context of HSCR therapy development.

**Figure 2:**
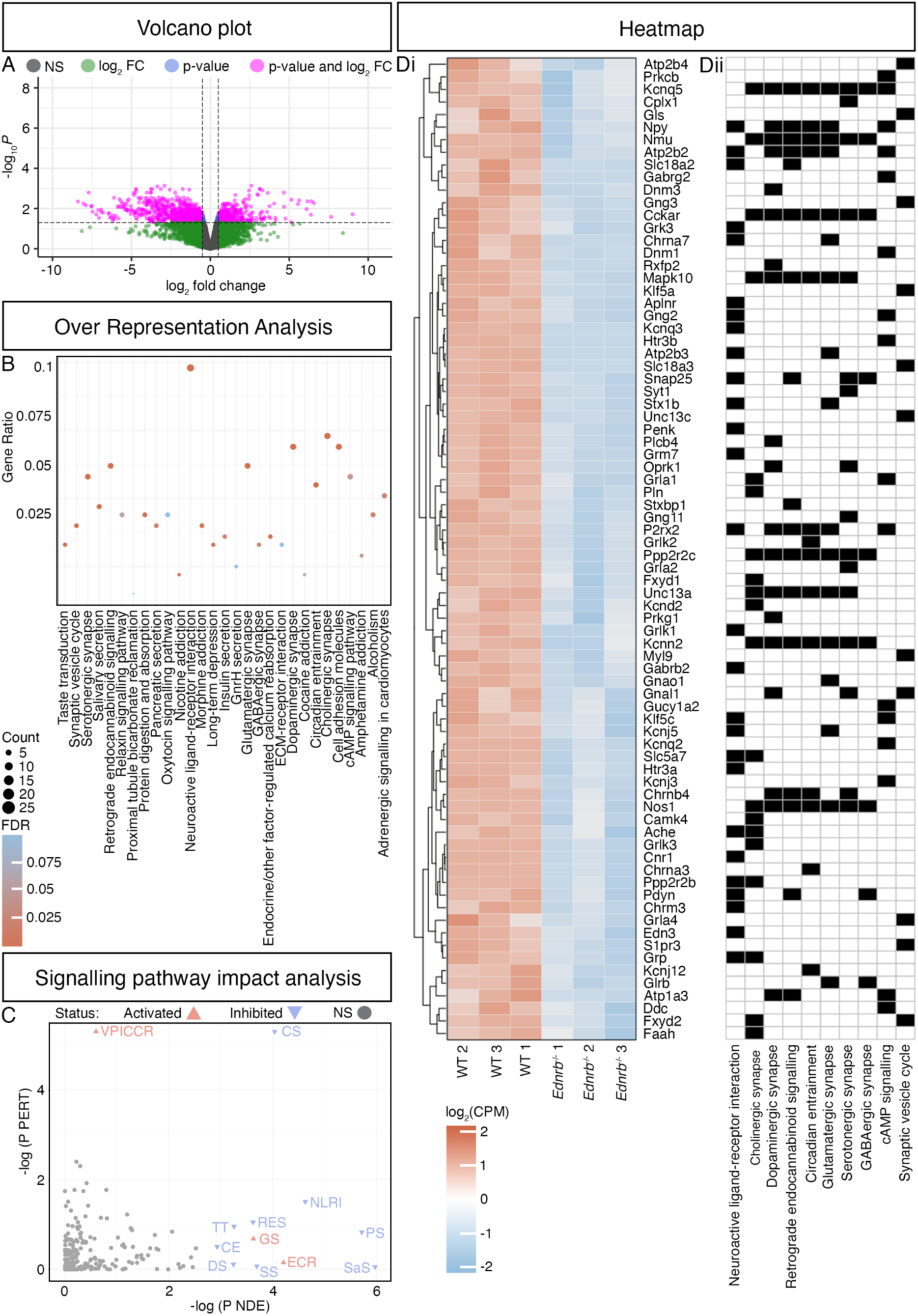
Expression and activity of numerous neuronal pathways are negatively impacted in B6;129-*Ednrb^tm1Ywa^/*J colonic tissue. (A) Expression patterns of RNA extracted from *Ednrb^-/-^* and WT littermates animals cluster by genotype, as assessed using multidimensional scaling (MDS), with 1,139 differentially expressed genes between *Ednrb^-/-^* and WT littermates. (B) Overrepresentation analysis showing KEGG pathways significantly enriched within the list of differentially expressed genes. Colour indicates magnitude of the fold discovery rate (FDR) and symbol size indicates the number of differentially expressed genes within each pathway. Gene ratio = fraction of down-regulated genes found in each pathway. (C) Signalling pathway impact analysis (SPIA) demonstrating the likely impact of the differentially expressed genes on the function of the pathway. NDE = number of differentially expressed genes, P PERT = probability of perturbation of the pathway. VPICCR – viral protein interaction with cytokine and cytokine receptor; CS – cholinergic synapse; NLRI – neuroactive ligand-receptor interaction; TT – taste transduction; RES – retrograde endocannabinoid signalling; PS – pancreatic secretion; CE – circadian entrainment; GS – glutamatergic synapse; DS – dopaminergic synapse; SS – serotonergic synapse; ECR – endocrine and other factor-regulated calcium reabsorption; SaS – salivary secretion. (Di) Heatmap showing all genes within the neuronal pathways significantly downregulated in *Ednrb^-/-^* animals compared to WT littermates. (Dii) Absence (white) or presence (black) of each gene in the affected pathways.

### Human ENS progenitors survive transplantation onto organotypically-cultured distal colons of *Ednrb^-/-^* mice and form ganglia-like structures

Previous work has demonstrated significant, positive increases in the contractile phenotype of human surgical discard HSCR samples following human ENS progenitor transplantation, with donor cells differentiating to appropriate neuroglial lineages (16). Having characterised the colonic pathology of *Ednrb^-/-^* animals, we wanted to determine whether transplantation of human vagal neural crest/early ENS progenitors, generated from hPSCs using our established protocol (15, 28), could restore the missing neuroglial cells in *Ednrb*^-/-^ aganglionic colonic tissue. Organotypically cultured colonic tissue isolated from *Ednrb^-/-^* mice were seeded with 500,000 ENS progenitors, derived from a hPSC line (SFCi55) marked by constitutive expression of the fluorescent reporter ZsGreen (29) and the transplanted tissue was maintained, *ex vivo*, for 21 days. Transplanted, ZsGreen+, cells condensed into discrete clusters connected by substantial TUBB3+ projections (**Fig. 3A**, arrows), with TUBB3+, ZsGreen+ projections extending between clusters (arrowheads) and into the host tissue (**Fig. 3A**, hollow arrowheads). Immunohistochemical examination of these structures revealed numerous TUBB3+, ZsGreen-labelled cells organised into ganglia-like structures (**Fig. 3B-G**, arrows) which often appeared alongside hypertrophied extrinsic innervation (**Fig. 3E-G**, hollow arrows). Further, we found that the dense clusters of ZsGreen+ transplanted cells contained instances of nitrergic and cholinergic differentiation, as evidenced by detection of nNOS+ (**Fig. 3H-J**, arrows) and VAChT+ neurons (**Fig. 3K-M**, arrows). These data confirm that hPSC-derived ENS progenitors survive transplantation into the aganglionic colon and have the capacity to differentiate to the major neuronal subtypes of the ENS in an aganglionic recipient microenvironment.

**Figure 3:**
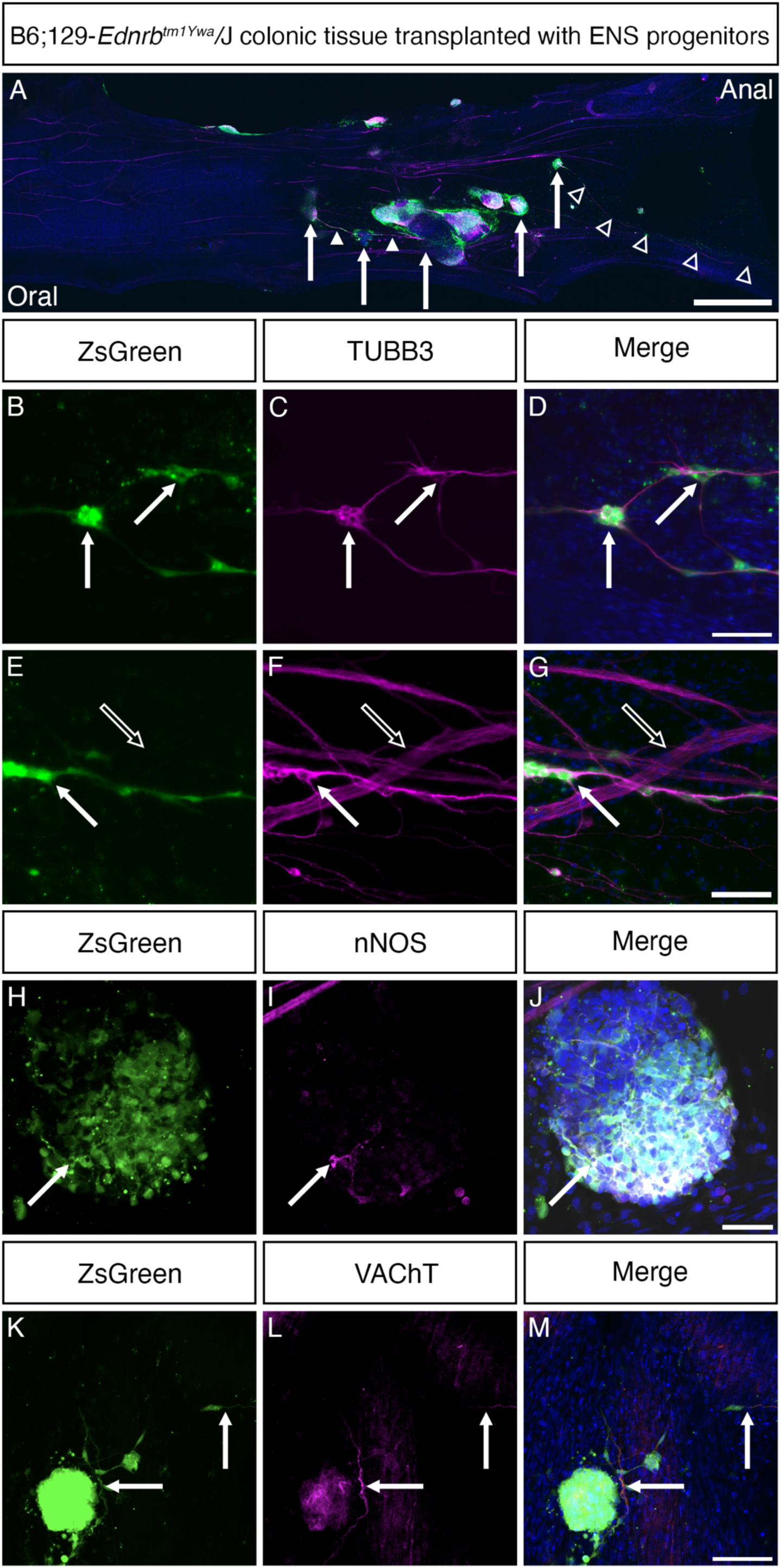
ZsGreen-labelled ENS progenitors survive transplantation and differentiate to form neuronal clusters in organotypically-cultured colonic tissue from B6;129-*Ednrb^tm1Ywa^*/J animals. (A) ENS progenitors survived 3 weeks (the latest time point examined) post-transplantation onto organotypically-cultured *Ednrb^-/-^* colonic tissue and formed dense clusters of TUBB3+, ZsGreen+ cells (A, arrows). These clusters extended projections both between clusters (arrowheads) and into the surrounding tissue (hollow arrowheads). (B-G) Confocal microscopy revealed the formation of ENS progenitor-derived ‘ganglia’-like interconnected structures (B-G, arrows). These structures appeared to interact with hypertrophied nerve bundles in the aganglionic region, presumably of extrinsic origin (E-G, hollow arrows). (H-M) Within ENS progenitor clusters we were able to detect nitrergic (H-J, arrows) and cholinergic neurons (K-M, arrows). Scale bars: A – 1mm, D, G - 100μm, J - 50μm, M - 100μm.

### Transplantation of organotypically*-*cultured *Ednrb^-/-^* distal colons with human ENS progenitors leads to significantly increased contractile activity

Having observed the integration of donor cells and the formation of ganglia-like structures composed of human ENS progenitor-derived neurons following transplantation, we next examined if the presence of donor cells exerted an effect on the contractile activity of previously aganglionic, *Ednrb^-/-^* distal colonic tissue. ENS progenitor-transplanted and sham-(vehicle-only) transplanted distal colonic tissue of *Ednrb^-/-^* mice were maintained *ex vivo* for 21 days. A section of transplanted tissue, approximately 1cm long, was collected and assessed using organ bath contractility (**Fig. 4A-B**). Compared to sham-transplanted tissue, ENS progenitor-transplanted tissue demonstrated significantly increased basal contractile frequency (3.38 +/-0.9784 contractions/min vs 1.07 +/- 0.309 contractions/min, p=0.0318, **Fig. 4C**). The basal contractile amplitude was not significantly affected in ENS progenitor-transplanted tissue compared to sham-transplanted tissue (37.78 +/- 5.829g and 22.83 +/- 6.72g, respectively, p=0.1177). Similarly, we detected no difference in the magnitude of basal contractions between sham-transplanted tissue and ENS progenitor-transplanted tissue (397.5 +/-185.1g.s and 193.3 +/-57.84g.s, respectively, p=0.3336).

**Figure 4:**
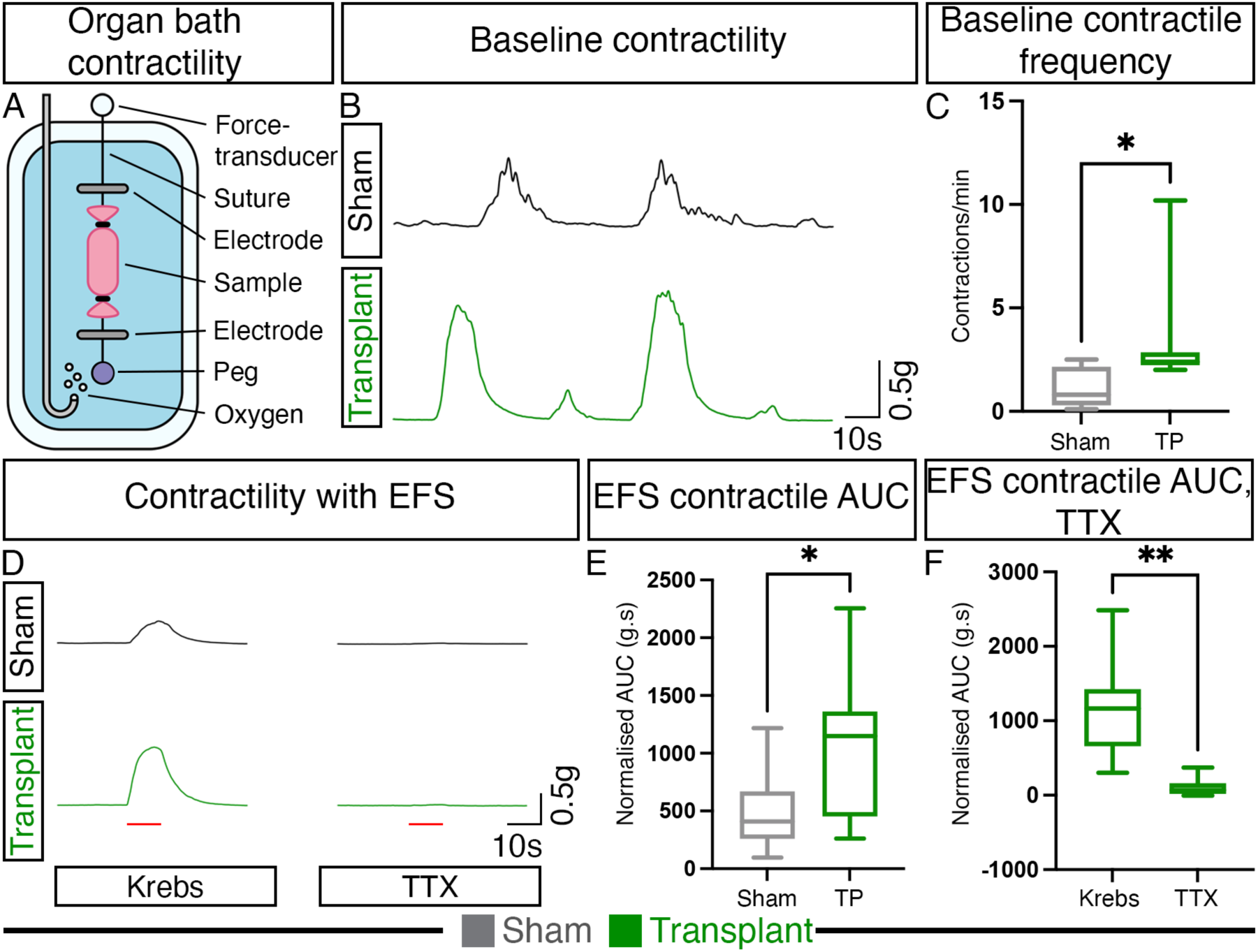
Transplantation of ENS progenitors onto organotypically-cultured colonic tissue from B6;129-*Ednrb^tm1Ywa^*/J animals results in significant contractile increases. (A) The effects of ENS progenitor transplantation on *Ednrb^-/-^* colonic tissue (denuded of the mucosa and cultured organotypically for 21 days) were assessed using organ bath contractility. (B) Representative traces of ENS progenitor-transplanted *Ednrb^-/-^*tissue (green trace) and sham-transplanted *Ednrb^-/-^* tissue (grey trace) samples at baseline (without stimulation). (C) Quantification confirmed a significantly increased contractile frequency in transplanted *Ednrb^-/-^*tissue. (D) Samples were subjected to electrical field stimulation (EFS) (for details, see methods). Red bar indicated timing and duration of EFS. (E) Transplanted samples shown increased contractile magnitude (area under the curve, or AUC). (F) Application of the sodium channel blocking agent TTX abolished contraction. Error bars are standard error of the mean. * indicates a p-value of <0.05; ** indicates a p-value of <0.01.

We next assessed the response of sham- and ENS progenitor-transplanted tissue to electrical field stimulation (EFS, **Fig. 4D**). Contractile magnitude, in response to EFS, was significantly increased in ENS progenitor-transplanted tissue compared to sham-transplanted tissue (1072.04.s and 487.04.s, respectively, p=0.0291, **Fig. 4E**). Further, in ENS progenitor-transplanted tissues, application of the sodium channel blocker TTX resulted in a significant decrease in contractile magnitude, following EFS (115.7 +/- 49.24g.s after TTX, 1167 +/- 232.1g.s before TTX, p=0.0011, **Fig. 4**F). These data suggest that transplantation and integration of hPSC-derived ENS progenitors leads to significantly increased, neuronally-mediated contractile responses within the aganglionic host tissue.

### Transplantation of human ENS progenitors partially rescues the biological pathways impacted in *Ednrb^-/-^* colonic tissue

To determine the impact of ENS progenitor transplantation on both the transplanted cells and the host microenvironment, we collected 6 additional *Ednrb^-/-^* mice and organotypically cultured the distal colonic tissue after either human ENS progenitor-transplant (n=3) or sham-(vehicle-only) transplant (n=3). Both groups were maintained organotypically for 21 days before RNA isolation. To enable comparison of human donor cells pre- and post-transplant, RNA was also isolated from 3 separate vials of freshly thawed human ENS progenitors (**Fig. 5A**). Our first RNAseq analysis focused only on human transcripts (i.e., exclusively donor-derived). PCA analysis revealed that post-transplant human donor cells clustered separately from control, non-transplanted “pre-transplant” ENS progenitors (**Fig. 5B**). Overrepresentation analysis revealed numerous significantly enriched pathways in post-transplanted human donor cells compared to an ENS progenitor state (**Tables S4, S5**). The majority of these suggested transplanted ENS progenitor-derived cells had undergone neuronal differentiation, with overrepresented pathways including *Synapse Organization* (FDR=3.42-20), *Neuron Projection Guidance* (FDR=1.46E-18) and *Axonogenesis* (FDR=1.2E-23, **Fig. 5C**, top 15 upregulated pathways are listed, see **Supp. Materials** for full list and **Fig. S3**). Within these pathways were key genes with well-documented importance in nervous system development, such as *Ret, Epha3*, *Epha4*, *Epha5*, *Epha6*, *Epha7*, *Robo2*, *Robo3, Slit1* and *Slit2*. Numerous WNT-associated genes involved in neural crest differentiation, including *Wnt3*, *Wnt3a*, *Wnt5a*, *Wnt7a*, *Wnt7b*, *Fzd3* and *Fzd9* were also significantly increased. Downregulated pathways included *Skeletal System Morphogenesis*, *Bone Morphogenesis* and *Blood Vessel Morphogenesi*s (**Fig. 5D**, top 15 downregulated pathways are listed, see **Table S5** for full list). Within these pathways were numerous genes associated with epithelial-mesenchymal transition and mesenchymal lineages, including *Twist1*, *Tbx3*, *Msx2*, *Gata3*, *Isl1*, *Myh9*, *Cnna* and *Bmp1/4,* potentially suggesting that the aganglionic gut microenvironment had biased donor cells toward neuronal differentiation at the expense of a mesenchymal fate. These findings suggest that the majority of donor human ENS progenitor cells have transitioned from an ENS progenitor/vagal NCC state toward neuronal differentiation within the host aganglionic tissue, supporting the conclusion that human ENS progenitors have the potential to repopulate a rudimentary ENS circuitry following transplantation into aganglionic colonic tissue.

**Figure 5:**
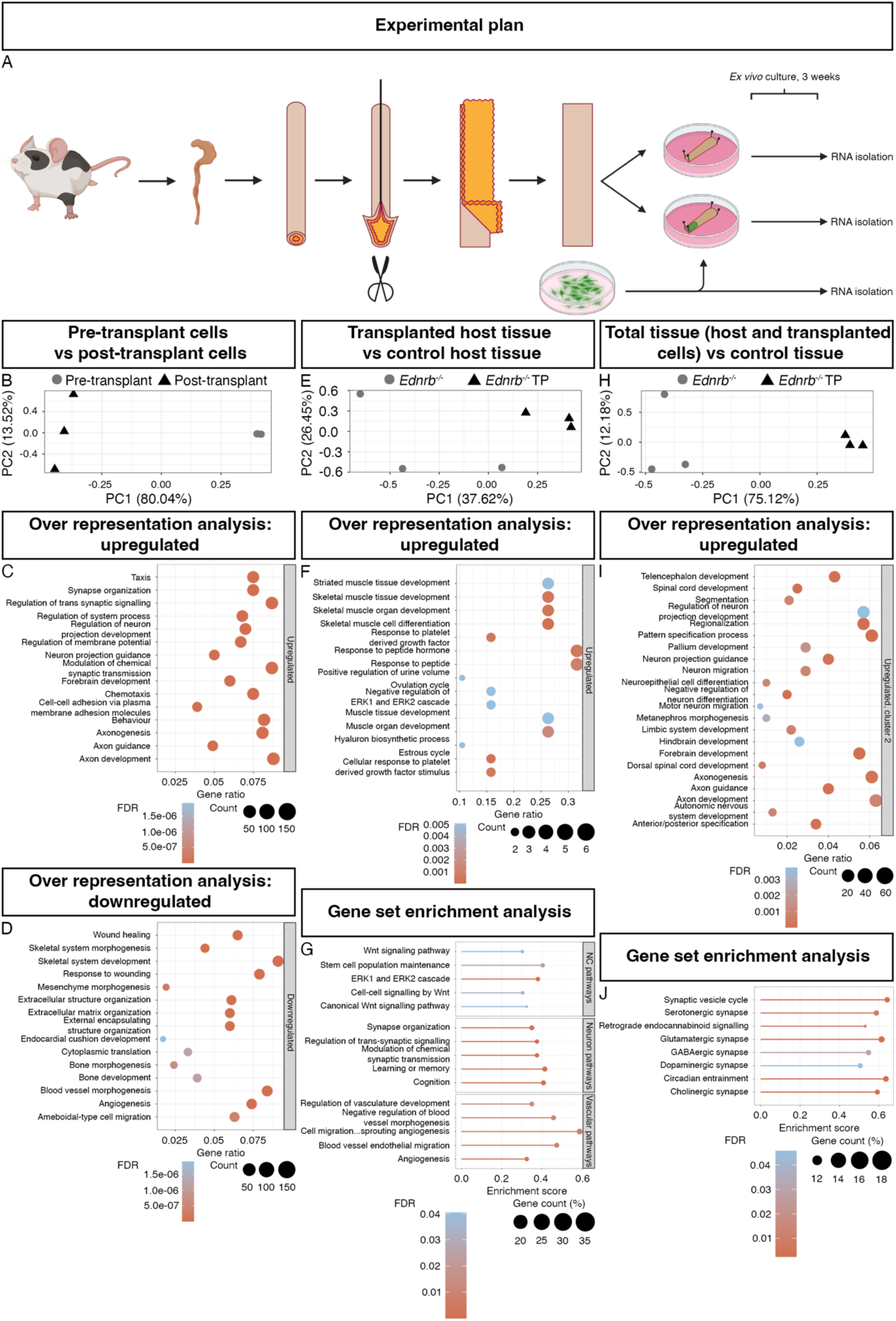
Transplantation of ENS progenitors onto organotypically-cultured colonic tissue from B6;129-*Ednrb^tm1Ywa^*/J animals rescues genetic pathways impacted in B6;129-*Ednrb^tm1Ywa^*/J colons. (A) Three experimental groups (n=3 for each group) were used for RNAseq analysis: (1) *Ednrb^-/-^* colonic tissue, denuded of the mucosa, sham-transplanted and cultured organotypically for 21 days; (2) *Ednrb^-/-^*colonic tissue, denuded of the mucosa, transplanted with ENS progenitors and cultured organotypically for 21 days; and (3) freshly thawed ENS progenitors. (B) Focussing on human-only (i.e., donor-derived) transcripts, principal component analysis (PCA) was used to assess clustering patterns of ENS progenitors pre-(grey circles) and post-(black triangles) transplant. (C) Overrepresentation analysis using the ‘Biological Pathways’ database revealed numerous pathways involved in neuronal differentiation (top 15 upregulated pathways are shown). (D) The top 15 downregulated pathways in post-transplant cells compared to pre-transplant. (E) Focussing on murine-only (i.e., host-tissue only) transcripts, PCA clustering revealed distinct clustering of sham-transplanted *Ednrb^-/-^* colonic tissue (grey circles) and ENS progenitor-transplanted *Ednrb^-/-^* colonic tissue (black triangles). (F) Overrepresentation analysis using the ‘Biological Pathways’ database revealed numerous pathways involved in muscle development. (G) Gene set enrichment analysis using the ‘Biological Pathways’ database revealed numerous significantly upregulated pathways, many of which seemed to fall into 3 categories: neural crest (NC) pathways; neuron pathways; and vascular pathways. (H) Considering both human and murine transcripts, PCA clustering revealed distinct clustering of sham-transplanted *Ednrb^-/-^* colonic tissue (grey circles) and ENS progenitor-transplanted *Ednrb^-/-^* colonic tissue (black triangles). (I) Overrepresentation analysis using the ‘Biological Pathways’ database revealed 4 unique clusters numerous based on expression patterns. Cluster 2 contained substantially greater numbers of affected pathways and so we focussed our analysis on this cluster (I), detecting numerous pathways related to neuronal development. (J) Gene set enrichment analysis (GSEA) focussing on the pathways previously found to have impacted activity in *Ednrb^-/-^*colonic tissue compared to *Ednrb^+/+^* colonic tissue, revealed significant upregulation of all pathways in ENS progenitor-transplanted *Ednrb^-/-^* colonic tissue compared to sham-transplanted colonic tissue. Colour indicates magnitude of the fold discovery rate (FDR). Symbol size indicates the number of differentially expressed genes within each pathway in overrepresentation analyses and the percentage difference in GSEA analyses. For full lists of all pathways see Supp. Materials.

To determine the molecular effect of ENS progenitor transplantation on host *Ednrb^-/-^* colonic tissue, we next focussed on murine transcripts only (i.e., exclusively recipient derived), ignoring donor cell-derived human transcripts. PCA analysis revealed that ENS progenitor-transplanted *Ednrb^-/-^* colonic tissue clustered separately from sham-transplanted *Ednrb^-/-^* tissue (**Fig. 5E, Table S6**). Overrepresentation analysis revealed multiple upregulated pathways in ENS progenitor-transplanted host tissue compared to sham-transplanted tissue, including *Striated Muscle Development* (FDR=5.0E-03), *Skeletal Muscle Cell Differentiation* (FDR=9.06E-06) and *Skeletal Muscle Tissue Development* (FDR=2.9E-04, **Fig. 5F**, top 15 upregulated pathways are listed, see **Table S7** for full list). Interestingly, one of the genes upregulated in these pathways was related to peripheral nervous system development: *Egr2*, potentially suggesting that the host microenvironment became more favourable to the development of a neo-ENS. Gene set enrichment analysis revealed significant increases in several key pathways, which we grouped into: (1) ‘neural crest (NC)-related pathways’, including *Wnt Signalling Pathway* (FDR=0.03) and *ERK1 and ERK2 Cascade* (FDR= 6.6E-04); (2) ‘neuron pathways’, including *Synapse Organization* (FDR=5.4E-04) and *Modulation of Chemical Synaptic Transmission* (FDR=6.9E-06); and (3) ‘vascular pathways’, including *Regulation of Vasculature Development* (FDR=0.01) and *Angiogenesis* (FDR=0.001, **Fig. 5G**, see **Table S8** for full list and **Fig. S3**). Notable genes with increased expression in these pathways include *Wnt2*, Wnt4, *Wnt11,* Wnt10a and *Wnt7b, Npy, Nrxn1*, *Asic2*, *Bdnf*, *Gdnf*, *Snca*, *Syn1*, *Snap25*, *Nos1*, *S100b* and *Kcn3*. To the best of our knowledge, this is the first time that transplantation of human ENS progenitors has been shown to result in substantial changes in the host molecular environment. Combined with the previous demonstration that transplanted human ENS progenitors overwhelmingly expressed gene pathways involved in neuronal differentiation and development, these data suggest that transplanted ENS progenitors are capable of both forming an ENS and ‘priming’ the aganglionic microenvironment to enable increased colonic motility.

Finally, we analysed both human and murine transcripts simultaneously to determine the overall effects of ENS progenitor transplantation into *Ednrb^-/-^* colonic tissue. PCA analysis revealed separate clustering of ENS progenitor-transplanted tissue and sham-transplanted tissue (**Fig. 5H, Table S9**). Overrepresentation analysis showed numerous upregulated pathways in ENS progenitor-transplanted tissue compared to sham-transplanted tissue, including *Regulation of Neuron Projection Development* (FDR=0.003), *Neuron Projection Guidance* (FDR=8.6E-05) and *Axon Development* (FDR=9.3E-04, **Fig. 5I**, top 22 upregulated pathways are listed, see **Table S10** for full list and **Fig. S3**). In our initial characterization of the transcripts affected by loss of *Ednrb* we had identified several pathways with significantly impacted activity compared to WT littermates (**Fig 2C**). To determine whether ENS progenitor transplantation had affected the expression these pathways we conducted gene set enrichment analysis and found that all pathways, which had been found to be downregulated in native *Ednrb^-/-^* colonic tissue, were significantly upregulated in ENS progenitor-transplanted tissue compared to sham transplanted tissue (**Fig. 5J, Table S11**).

Taken together, these data suggest that transplantation of ENS progenitors to aganglionic colonic tissue drives transplanted human ENS progenitor cells towards a neuronal fate and simultaneously adapts the aganglionic host microenvironment both for increased contractile motility and survival of neural crest-derived cells (for example, via increased expression of Wnt/Erk pathways). Combined, these have the effect of at least partially rescuing all impacted neuronal pathways in the *Ednrb^-/-^* aganglionic colon, demonstrating the potential of human ENS progenitors to restore multiple biological pathways impacted in HSCR.

## Discussion

Over the past 15 years, significant progress has been made in the development and pre-clinical application of regenerative medicine approaches utilising cell-based therapies to replace the ENS missing in HSCR (16, 30, 31). Numerous studies have shown that transplanted ENS progenitors/stem cells (both murine and human derived) can integrate with host tissue (20, 32, 33). Of note, many of these studies utilise animal models recapitulating human HSCR pathology, including mouse lines targeting the *Ednrb* gene (18, 20, 34). However, significant variability exists between the various *Ednrb* mouse lines which may limit their applicability for comparative studies. Additionally, recent work has demonstrated the substantial impact that lab-to-lab variability can have on the phenotype of a given model, including the specific diet the mice are maintained on, strongly advocating in-house characterization of all mouse lines prior to their use (35). In the current manuscript, we undertook an in-depth characterization of B6;129-*Ednrb^tm1Ywa^*/J mice, assessing the anatomical, functional and molecular deficits as a baseline before application of ENS progenitors. Following transplantation into aganglionic colonic tissue isolated from *Ednrb^-/-^* mice, we demonstrated (i) the integration and neuronal differentiation of human ENS progenitor-derived cells, (ii) an increased contractile phenotype and (iii) restoration of disease-impacted pathways, with progenitor-induced adaptation of the host microenvironment towards a neural crest-supportive state.

We have previously shown significant increases in the contractile behaviour of human surgical discard tissue taken from the aganglionic colon of HSCR patients following transplantation with human hPSC-derived vagal neural crest-ENS progenitors (16). Here, we show similar results following transplantation of human ENS progenitors (generated using the same protocol) into organotypically cultured aganglionic *muscularis* tissue of *Ednrb^-/-^* mice. Notably, the functional observations we present here reflect similar increases in contractile activity observed following the transplantation of allogenic murine enteric neural stem cells in the same B6;129-*Ednrb^tm1Ywa^*/J model (16). Hence, the reproducibility of our observations of significantly increased basal contractile frequency (including response to EFS and sensitivity to TTX) suggests that our *ex vivo* transplantation methodology provides a tractable platform to study the integration and effects of human ENS progenitor cell transplantation.

While previous studies have highlighted the capacity of donor ENS cell types from various sources to integrate within mouse models of HSCR, our approach provides vital data on the molecular consequences of ENS progenitor transplantation. Using native *Ednrb^-/-^* and wildtype littermate tissue, we highlight differences in gene expression and biological pathways, as a consequence of *Ednrb* knockout in the B6;129-*Ednrb^tm1Ywa^*/J model. Unsurprisingly, given the severe aganglionic phenotype observed in this model, the majority of these were neuronal pathways, including the *Serotonergic synapse*, *Glutamatergic synapse*, *GABAergic synapse*, *Dopaminergic synapse*, *Cholinergic synapse* pathway and *Neuroactive ligand-receptor interaction* pathways. Following *ex vivo* transplantation, our approach (i.e., human ENS progenitors transplanted to murine *Ednrb^-/-^*colon) enabled the separation and analysis of human and mouse transcripts. Critically, this methodology provided novel data on both cell autonomous effects, in donor-derived cells, and non-cell autonomous effects within the recipient microenvironment. Numerous neuronal differentiation pathways were found to be upregulated in ENS progenitor-derived human cells post-transplant compared to control “pre-transplant” ENS progenitors, suggesting their potential to generate ENS cell types in aganglionic tissue. This was further supported by immunohistochemical detection of neuronal subtype protein expression in donor-derived cells in transplanted tissue. Downregulated pathways in donor-derived human transcripts compared to “pre-transplant” ENS progenitor cells included mostly ‘morphogenic’ pathways, suggesting a shift towards a more mature cell phenotype, further supporting differentiation of ENS progenitor-derived cells post-transplant. GSEA of murine-only transcripts, as a proxy for the host microenvironment, revealed numerous significantly increased pathways relating to vascularization, muscle development and neural crest-related pathways, potentially creating a permissive environment for ganglia formation and restoration of colonic motility. Finally, analysis of human and murine transcripts simultaneously revealed that all pathways impacted in our native tissue analysis (i.e., *Ednrb^-/-^*colon compared to WT littermate tissue), as revealed by SPIA during model characterization, had been significantly increased following ENS progenitor transplantation.

Taken together, these results suggest that following transplantation, human ENS progenitors undergo differentiation towards a neuronal phenotype and that ENS progenitor transplantation may have wide ranging effects, rescuing many of the molecular features of aganglionosis.

In conclusion, our findings confirm the validity of the B6;129-*Ednrb^tm1Ywa^*/J mouse model as a tool for testing novel HSCR therapies. Furthermore, our data suggest that human ENS progenitor transplantation may have wide ranging, positive effects, including (i) donor-derived, ganglia-like structures comprised of neuroglial cells; (ii) increased contractile function and (iii) at least partial rescue of the molecular pathogenesis found within aganglionic tissue and significant upregulation of ENS-permissive pathways. These data provide strong support for the translational development of ENS progenitors.

## Materials and Methods

### Animals

B6;129-*Ednrb^tm1Ywa^*/J mice were obtained from Jackson laboratories (Bar Harbor, MN, USA) and maintained with heterozygous (*Ednrb^+/-^*) breeders. Offspring of heterozygous breeding pairs were genotyped to confirm homozygous loss of the *Ednrb* gene (*Ednrb^-/-^*). Homozygous WT (*Ednrb^+/+^*) littermates were used as controls. Animals were maintained and used for experiments in accordance with the UK Animals (Scientific Procedures) Act 1986 following approval by the University College London Biological Services Ethical Review Process. Animals were maintained on 2018 Teklad Global 18% Protein Rodent Diet (Inotiv, UK). For microbial status, see **Table S13.** Animal husbandry at UCL Biological Services was in strict accordance with the UK Home Office Certificate of Designation.

### Immunofluorescence

Tissue samples were fixed in paraformaldehyde (4% w/v) at RT for 1 hour and stored in PBS. Samples were blocked with 1% bovine serum albumin (Sigma Aldrich, UK) and 0.1% Triton X-100 (Sigma Aldrich, UK) in 1XPBS at RT before incubation in primary antibody (**Table 1**) diluted in blocking solution overnight at 4°C. Secondary antibodies (**Table 1**) diluted in blocking solution were applied for 1 hour at RT. Slides were cover-slipped using Vectashield (Hard Set, Dako, UK). Slides were stored at 4°C until imaging.

**Table 1:**
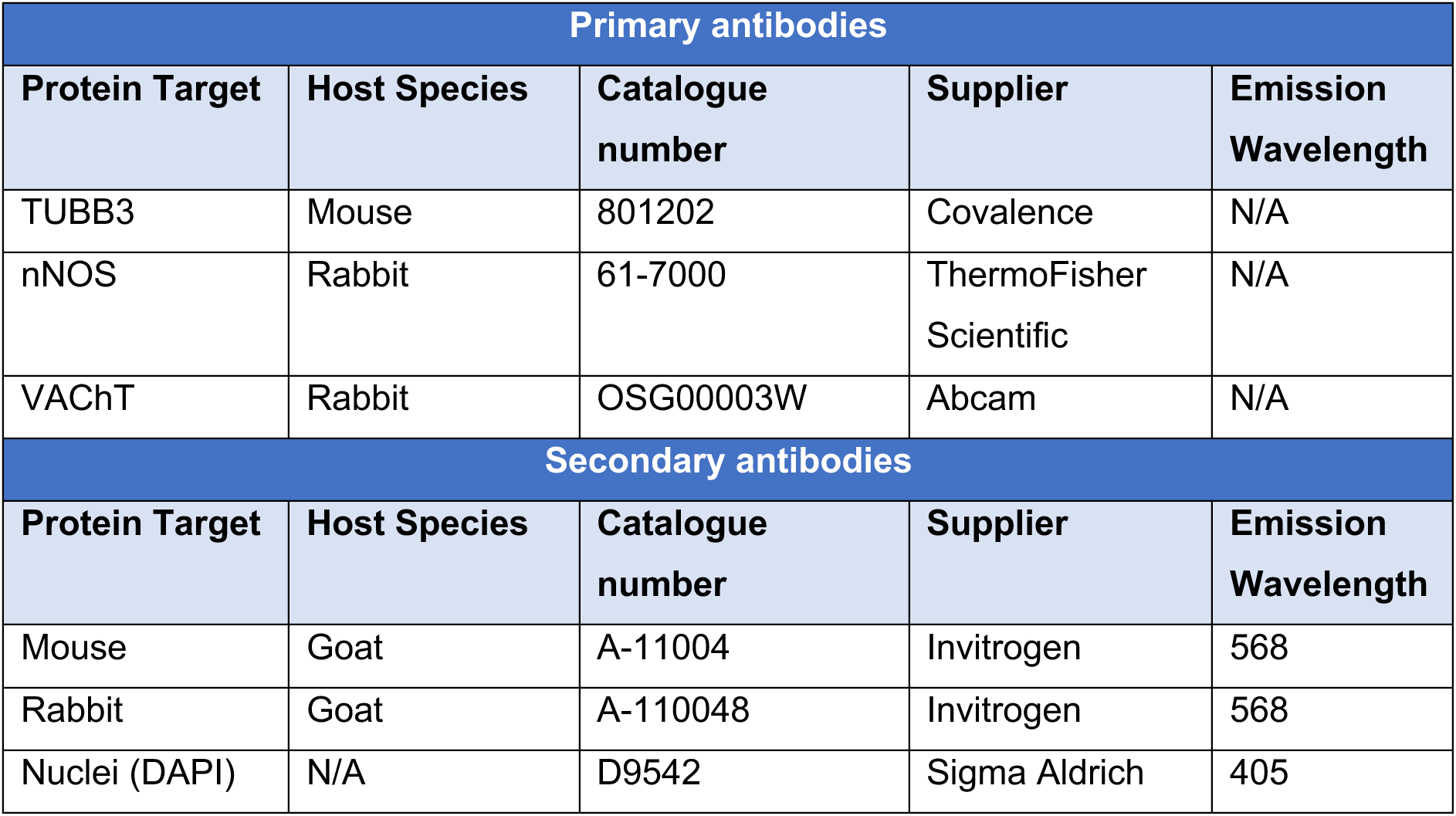
Antibodies used for immunofluorescence investigations.

| Primary antibodies |  |  |  |  |
| --- | --- | --- | --- | --- |
| Protein Target | Host Species | Catalogue number | Supplier | Emission Wavelength |
| TUBB3 | Mouse | 801202 | Covalence | N/A |
| nNOS | Rabbit | 61-7000 | ThermoFisher Scientific | N/A |
| VACHT | Rabbit | OSG00003W | Abcam | N/A |
| Secondary antibodies |  |  |  |  |
| Protein Target | Host Species | Catalogue number | Supplier | Emission Wavelength |
| Mouse | Goat | A-11004 | Invitrogen | 568 |
| Rabbit | Goat | A-110048 | Invitrogen | 568 |
| Nuclei (DAPI) | N/A | D9542 | Sigma Aldrich | 405 |

### RNA extraction

RNA was extracted from colonic muscularis tissue isolated from *Ednrb^-/-^*mice and wild type littermates. Colonic tissue was isolated, opened along the oro-anal axis and denuded of the mucosa using fine forceps. Additional RNA was isolated from colonic muscularis tissue isolated from *Ednrb^-/-^* animals transplanted with either ENS progenitors or vehicle only (sham) and organotypically-cultured for 3 weeks. For both experiments, the distal third of the colonic muscularis was collected and snap frozen in liquid nitrogen until needed. Tissue was homogenised using TissueLyser 5mm Beads (Qiagen, UK) with moderate vortexing and total RNA extracted using an RNeasy mini kit (Qiagen, UK) according to the manufacturer’s instructions, including an optional treatment with DNase I (Qiagen, UK).

### mRNA Library preparation

Total RNA quantification and integrity was confirmed using Agilent’s 4200 Tapestation (Standard Total RNA assay). RINs values were confirmed to all be > 7.0, indicating high integrity RNA suitable for library prep. For each sample, 500ng of total RNA were processed using the KAPA mRNA HyperPrep Kit (Roche p/n KK8580) according to manufacturer’s instructions. Briefly, mRNA was isolated from total RNA by use of paramagnetic Oligo dT beads to pull down poly-adenylated transcripts. The purified mRNA was fragmented using chemical hydrolysis (heat and divalent metal cation) and primed with random hexamers. Strand-specific first strand cDNA was generated using Reverse Transcriptase in the presence of Actinomycin D. The second cDNA strand was synthesised using dUTP in place of dTTP, to mark the second strand. The resultant cDNA was “A-tailed” at the 3’ end to prevent self-ligation and adapter dimerisation. Full length xGen adaptors (IDT), containing two unique 8bp sample specific indexes, a unique molecular identifier (N8) and a T overhang were ligated to the A-Tailed cDNA. Successfully ligated cDNA molecules were enriched with limited cycle PCR (12 PCR cycles).

### mRNA sequencing

High yield, adaptor-dimer free libraries were confirmed on the Agilent TapeStation 4200 (High Sensitivity Agilent DNA 1000 assay). Samples were quantified using the Qubit High Sensitivity DNA assay and normalised to 10nM. An equal volume of each library was pooled together and re-quantified by Qubit. Samples were sequenced on the NextSeq 2000 instrument (Illumina, San Diego, US) at 800pM, using a 57bp paired-read read run with corresponding 8bp dual sample indexes and 8bp unique molecular index.

### Sequencing Data Analysis

BCL data were demultiplexed and converted to fastq files using Illumina’s *BCL Convert* Software v3.7.5. During this process, Unique Molecular Index (UMI) sequence from the IDT xGen UMI-UDI adapter were removed from the i7 sequence and added into the fastq read header. Resultant fastq files were then adapter and quality trimmed (*<u>fastp</u>*v0.20.1) before being aligned to the mouse genome (UCSC mm10) using *RNA-STAR* 2.5.2b. Aligned reads were UMI deduplicated using Je-suite (1.2.1). QC of alignment was performed by *<u>Picard</u> CollectRNASeqMetrics* and, if paired-end data, *CollectInsertSizeMetrics*. All fastp, RNA-STAR and Picard metrics were collated into a primary *<u>MultiQC</u>* report. The deduplicated alignments were used to generate reads per transcript with *<u>FeatureCounts</u>* along with appropriate GTF annotation. All annotation and sequences were obtained from Illumina <u>iGenomes</u>.

### RNA-seq analysis

The count table obtained with FeatureCounts was imported into R/v4.1.3 (36). The transcriptome was filtered by using the function “filterByExpr” of the edgeR package (37) (with parameters “min.count=5; min.total.count=15; min.prop=0.7”) to remove genes with low (or null) counts that would contribute to background noise (38) and was then normalised by using the TMM normalisation (39).

With the normalised counts, an initial unsupervised analysis was performed by applying the multi-dimensional scaling (MDS) algorithm of the edgeR package to assess the clustering of samples in a 2-dimensional space. Then a formal pairwise differential expression analysis was performed using the edgeR package (genes with FDR < 0.05 and Fold Change ≥ |1.5| were deemed significant).

To determine if our lists of up- and down-regulated genes were enriched more than would be expected for genes belonging to a specific pathway, an over-representation analysis was run with the package clusterProfiler (40) by using the KEGG database and comparing our lists of differentially expressed genes against the filtered transcriptome, as suggested by (41). Pathways with FDR < 0.1 were deemed significant.

A signalling pathway impact analysis (SPIA) was finally performed with the package SPIA (27) on the entire list of differentially expressed genes with their fold change information. This analysis has the advantage of imposing on the classical over-representation analysis, the complex gene interactions that characterise molecular pathways, to determine if they are impacted or not by the changes in gene expression. Pathways with a global FDR < 0.1 were deemed significant.

Plots were performed using the ggplot2 (42), complexHeatmap (43) and EnhancedVolcano libraries.

For the transplant experiment, fastq files that contained both human and mouse were initially split to separate human reads from mouse reads. This step was achieved via the byPrim.py script originally created by Song and colleagues (44) and refined by Chadarevian et al. (45). This protocol involves the creation of a hybrid genome joining human and mouse genomes and aligning the reads on the hybrid genome to sort them by their primary alignment in separate fastq files. Results of the separation are reported in **Table S12** and **Fig. S3A**. After read separation, mapping with STAR was carried out on each respective original genome (i.e., mm10 for mouse and hg38 for human reads) and subsequent counts were generated with FeatureCounts (with options -M and --fraction). Count tables were imported in R and analysed as described previously.

To determine the overall effects of ENS progenitor transplantation into *Ednrb^-/-^* colonic tissue, we analysed both human and murine transcripts simultaneously. The program *Orthofinder* (46) was used to define orthogroups between the two species; subsequently each gene ID in the count tables was replaced with its orthogroup ID and the two count tables were merged, summing the counts for the genes belonging to the same orthogroup. The resulting count table was imported in R and analysed as described previously.

### Processing of mouse tissue for *ex vivo* culture

*Ednrb^-/-^* animals and WT littermates were collected at P14 and killed by cervical dislocation. The entire intestinal tract was harvested into sterile Ca^2+^/Mg^2+^ free phosphate buffered saline (PBS, 0.1mol L-1, pH 7.2, Gibco Life Technologies, UK). The majority of the mesentery was removed via fine dissection and the colon isolated and opened along the oral-anal axis at the mesenteric border. Colonic tissue was stretched and pinned onto Syglard-based dishes and denuded of the mucosa. Samples were transferred to fresh Syglard-based dishes and pinned using tungsten wire (0.05mm, Thermo Scientific, UK) with the serosal side facing upwards in DMEM/F12 media (Gibco Life Technologies, UK) media with L-Glutamine and HEPES, supplemented with primocin (500mg/mL, Invivogen, UK), N2 (Gibco Life Technologies, UK), and B27 (Gibco Life Technologies, UK). Following washes with several media changes, samples were placed in a humidified incubator (37°C, 5% CO2), overnight if the tissue was to receive transplants of either ENS progenitors or vehicle only, or else for 3 weeks, with media changes every other day.

### Cells

Human pluripotent stem cell (PSC)-derived vagal neural crest cells (ENS progenitors) were generated as previously described (15, 28). The ZsGreen-labelled hPSC line SFCi55-ZsGr was used. Briefly, cells were seeded onto Geltrex-coated plates in the presence of CHIR 99021 (Tocris, UK), SB 431542 (Tocris, UK), DMH-1 (Tocris, UK), BMP4 (Fisher Scientific, UK) and Rho-associated coil kinase (ROCK) inhibitor Y-27632 2HCl (Generon, UK), with media replenished on day 2 excluding ROCK inhibitor. After 4 days, media was supplemented with retinoic acid (Merck, UK) for a further 2 days. Cells were collected and cryopreserved until required.

### Transplanting organotypically-cultured B6;129-*Ednrb^tm1Ywa^*/J colonic tissue with human ENS progenitors

On the day of transplantation, frozen cell aliquots were thawed and resuspended (125,000 cells/μL) in culture media (DMEM F12 media with L-Glutamine and HEPES, supplemented with primocin, N2 and B27. Cell viability was confirmed using trypan blue dye (Fisher Scientific, UK) and an automated cell counter (TC20 Automated Cell Counter, Biorad, UK). Organotypically-cultured *Ednrb^-/-^*colonic muscularis tissue samples were drained of media, removing as much liquid as possible from the edges of the tissue. 5μL of cell suspension was pipetted onto the anal-most portion of the tissue (500,000 cells total). After 30 minutes media was gently replaced and the samples returned to the incubator. Sham transplants underwent the same procedure, with a transplant of 5μL vehicle-only. Transplanted and sham-transplanted organotypically-cultured samples were maintained for 3 weeks, with media replaced every other day.

### Contractility assays

Organotypically-cultured colonic tissue samples from B6;129-*Ednrb^tm1Ywa^*/J mice were transferred to oxygenated Krebs (NaCl 120.4mM, KCl 5.9mM, NaHCO_3_ 15.5mM, glucose 46mM, MgCl_2_ 10.3mM, NaH_2_PO_4_ 6.2mM, CaCl^2^ 3.3mM) solution and the anal-most segment (approximately 1cm in length) isolated for contractility experiments. Each segment was folded in half along the oral-anal axis and mounted in organ baths (10 ml, SI-MB4; World Precision Instruments Ltd, UK) connected to force transducers (SI-KG20, World Precision Instruments Ltd, UK) using fine suture (size 4.0, Fine Science Tools, Germany) under an initial tension of 0.5g. An organ bath temperature of 37°C was maintained with perfusion of fresh, oxygenated Krebs solution. Following 1 hour of equilibration, a Lab-Trax-4 data acquisition system (World Precision Instruments Ltd, UK) was used to record contractile activity. Tissue samples were subjected to electrical field stimulation (EFS) for 30s (5Hz; 40V; 0.3ms pulse duration) at intervals of 60s via platinum electrode loops located at both ends of the tissue sample using a MultiStim System (D330, World Precision Instruments Ltd, UK), in the presence and absence of TTX (1µM, Cambridge Bioscience, UK).

Contractility data were collected, stored and analysed via Labscribe 2-software (World Precision Instruments Ltd, UK). Baseline contractile frequency was determined by counting the number of contractions over a 20-minute period. The baseline contractile amplitude was determined by measuring difference between the baseline tension and the peak of each contraction (g). The contractile response to electrical stimulation was calculated by measuring the area under the curve (AUC) for the duration of the electrical stimulation, referred to as baseline stim AUC.

### Statistics

Data was statistically analysed and graphs generated using GraphPad Software (GraphPad Prism). All data are expressed as mean ± standard error of the mean. The significance of functional data was assessed using a paired Student’s t-test, with p-values <0.05 taken as significant. The significance of Mendelian inheritance adherence and gender bias was assessed using a Chi squared test, with p-values <0.05 taken as significant.

## Acknowledgements

The authors thank Dr Dale Moulding (UCL Great Ormond Street Institute of Child Health Imaging Facility) for technical assistance. We acknowledge Anthony Brooks at UCL Genomics for carrying out the library preparation, sequencing, and analysis. Part of this research was conducted at UCL Great Ormond Street Institute of Child Health and supported by the NIHR Great Ormond Street Hospital Biomedical Research Centre. Views expressed in this manuscript are solely those of the authors and not necessarily those of the NHS, the NIHR or the Department of Health.

## Author contributions

B.J., F.C., L.P., S.P., S.L. and T.M. acquired and interpreted data. B.J., F.C., P.D.C., P.W.A., A.T. and C.J.M. contributed to study concept and design and interpreted data. A.T., C.J.M. acquired funding. B.J., F.C., L.P., S.P., P.W.A., A.T. and C.J.M. drafted the manuscript.

## Marterials and Correspondence

Materials and data can be provided by the corresponding author pending scientific review and a completed material transfer agreement

## Transcript profiling

All sequencing data have been deposited on NCBI’s SRA. Further, data have been deposited at GEO. All data will be made available following the cessation of the embargo.

## Data availability

All data needed to evaluate the conclusions in the paper are present in the paper and/or the Supplementary Materials. All sequencing data have been deposited on NCBI’s SRA. Further, data have been deposited at GEO. All data will be made available following the cessation of the embargo.

