## Supplementary figures 1, 2 and 3 for "Functional and molecular rescue of aganglionic colon by human enteric nervous system progenitor transplantation in Hirschsprung disease"

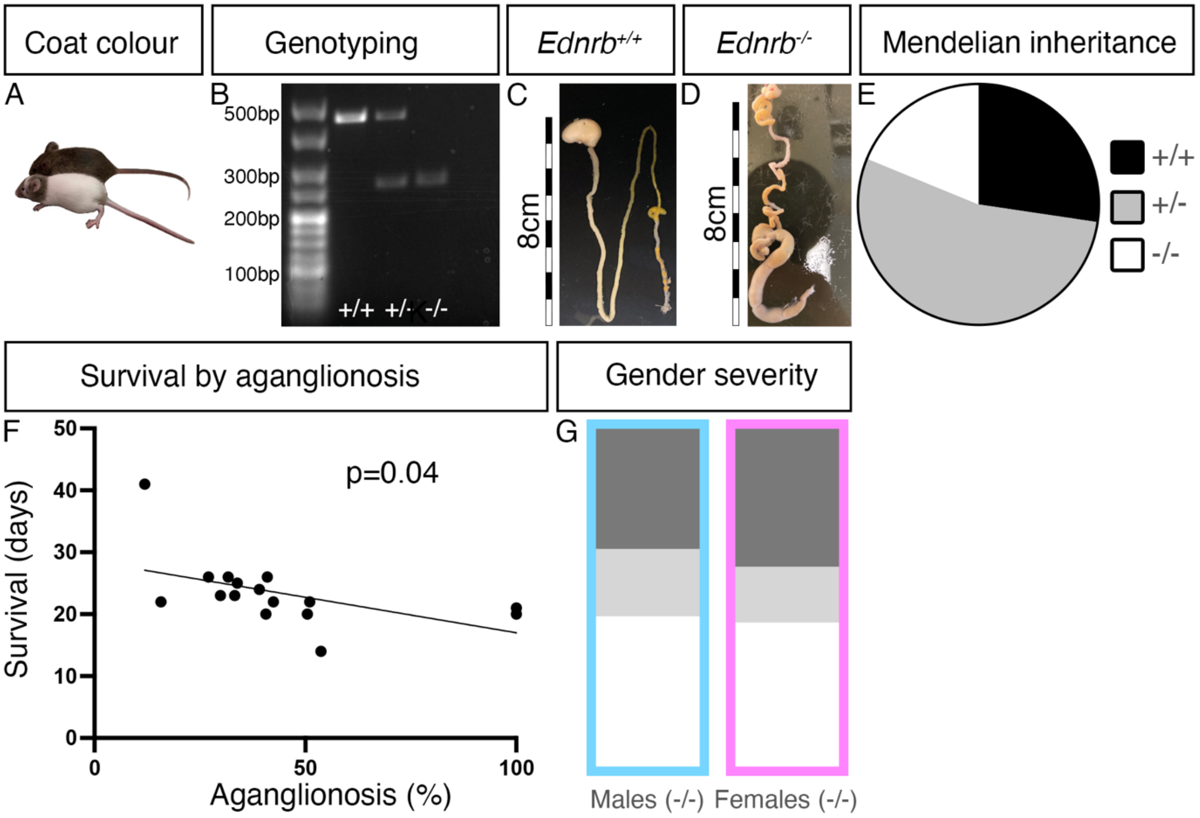


**Fig S1: B6;129-*Ednrb*$^{tm1Ywa}$/J animals can be easily identified and phenocopy pathology observed in human Hirschsprung disease**. (A) *Ednrb*$^{-/-}$ animals display characteristic skewbald coat colour. (B) PCR analysis confirms *Ednrb*$^{-/-}$ genotype. (C) *Ednrb*$^{+/+}$ wild type littermates display expected colon morphology, including well defined, regular pellets. (D) Conversely, *Ednrb*$^{-/-}$ animals displayed constricted distal colons and swollen megacolon. (E) Assessment of the proportion of *Ednrb*$^{-/-}$, *Ednrb*$^{+/+}$ and *Ednrb*$^{+/-}$ pups in 13 litters (124 pups total). (F) The degree of aganglionosis was variable, with severity significantly associated with reduced survival. (G) Aganglionosis severity did not differ between the genders (length of ganglionic tissue: dark grey; length of transition zone tissue: light grey; length of aganglionic tissue: white).

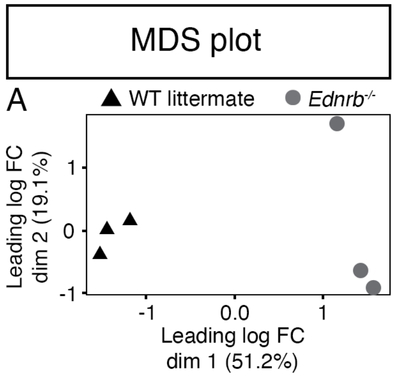


**Figure S2: Bulk RNAseq reveals distinct gene expression patterns in *Ednrb*$^{-/-}$ and WT littermate distal colonic tissue.** (A) Principal component analysis reveals separate clustering of *Ednrb*$^{-/-}$ aganglionic muscularis tissue and muscularis tissue derived from wild type littermates.

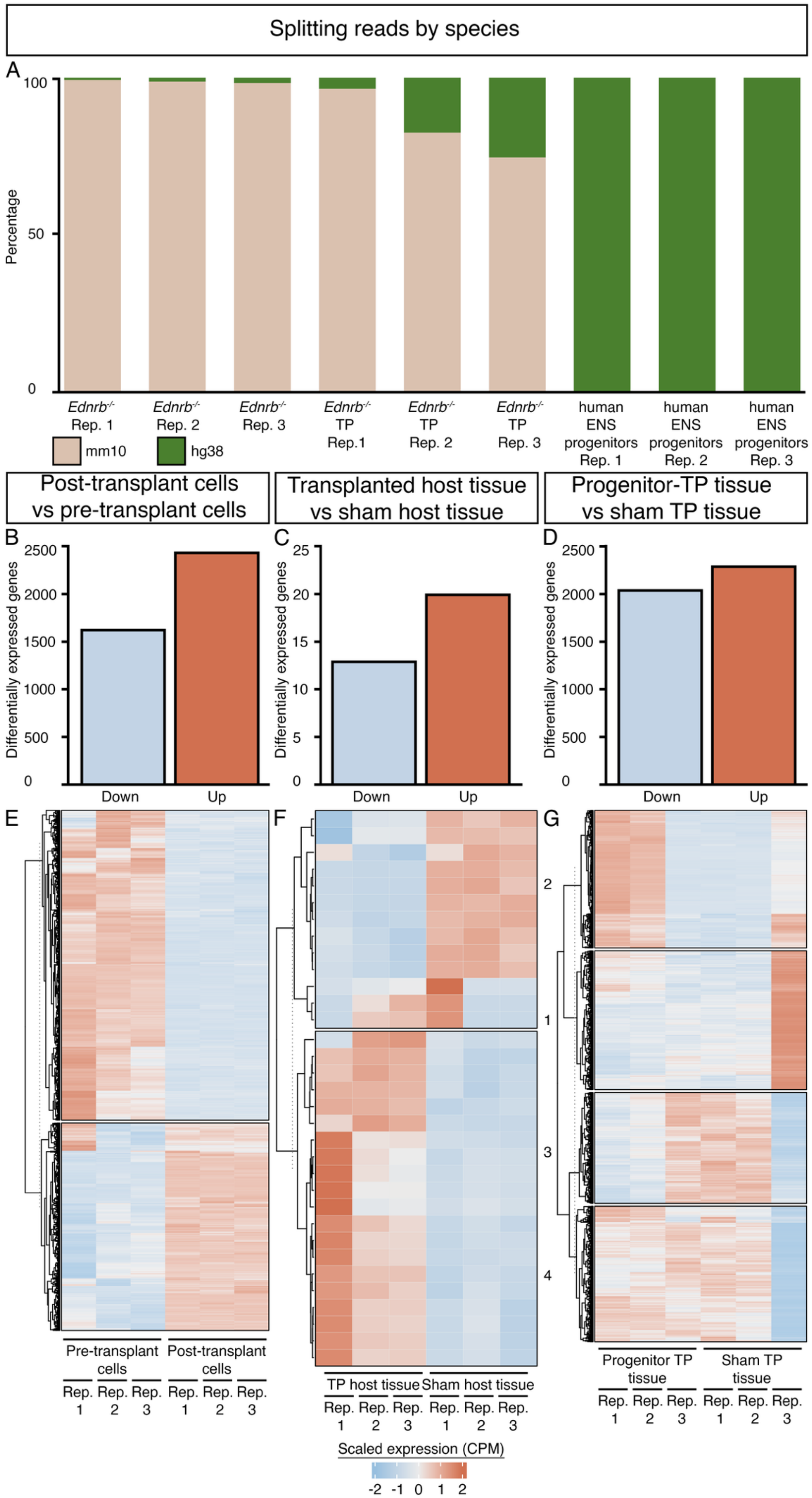

Splitting reads by species
A
100
50
0
Percentage
Ednrb-/- Rep. 1
Ednrb-/- Rep. 2
Ednrb-/- Rep. 3
Ednrb-/- TP Rep.1
Ednrb-/- TP Rep. 2
Ednrb-/- TP Rep. 3
human ENS progenitors Rep. 1
human ENS progenitors Rep. 2
human ENS progenitors Rep. 3
mm10
hg38
Post-transplant cells vs pre-transplant cells
Transplanted host tissue vs sham host tissue
Progenitor-TP tissue vs sham TP tissue
B
C
D
Differentially expressed genes
2500
2000
1500
1000
500
0
25
20
15
10
5
Down
Up
E
F
G
2
1
3
4
Pre-transplant cells
Post-transplant cells
TP host tissue
Sham host tissue
Progenitor TP tissue
Sham TP tissue
Rep. 1
Rep. 2
Rep. 3
Scaled expression (CPM)
-2 -1 0 1 2

**Figure S3: Species-specific transcripts from combined bulk RNAseq can isolated for individual analysis.** (A) Human (HG38, green) and murine (mm10, beige) transcripts could be isolated using the protocol established by Song *et al* and refined by Chadarevian *et al* (44, 45). (B) Considering human-only transcripts (i.e., donor-derived) 1641 genes were downregulated and 2447 genes were upregulated in transplanted human ENS progenitors compared to pre-transplant human ENS progenitors. (C) Considering murine-only transcripts (i.e., host tissue-derived), 13 genes were down-regulated and 20 genes were upregulated in host tissue transplanted with human ENS progenitors compared to host tissue receiving a sham transplant. (D) Considering both human and mouse transcripts simultaneously, 2053 genes were down-regulated and 2302 genes were upregulated in human ENS progenitor-transplanted tissue compared to sham-transplanted tissue. (E) Heatmap showing up-and down-regulated genes in post-transplanted human ENS progenitors compared to pre-transplant human ENS progenitors (human-only genes, see **Supp. Table 5** for full list). (F) Heatmap showing up-and down-regulated genes in host tissue transplanted with human ENS progenitors compared to host tissue receiving a sham transplant (murine-only genes, see **Supp. Table 8** for full list). (G) Heatmap showing up-and down-regulated genes in host tissue transplanted with human ENS progenitors compared to host tissue receiving a sham transplant (both human and murine genes, see **Supp. Table 10** for full list). Up- and down-regulated genes were clustered according to Rep. = replicate, TP = transplanted.
